# SICM6A: Identifying m^6^A Site across Species by Transposed GRU Network

**DOI:** 10.1101/694158

**Authors:** Wenzhong Liu

## Abstract

N6-methyladenosine (m^6^A) is the most prevalent cross-species RNA methylation modification and plays a pivotal role in various biological processes. The biochemical methods to find m^6^A sites are expensive and time-consuming, and the false positive rate of identified sites is high relatively. Meanwhile, the current computations are complex, and the prediction performance is relatively low both on little data sets and large data sets. This paper, at this point, presents a deep learning model with a transposed operation in the middle of GRU layers, SICM6A, for identifying m^6^A sites across-species. It adopts the mixed precision training manner to improve the speed and performance, and predicts m^6^A sites only by directly reading the *3-mer* encoding of the m^6^A short sequence. The cross-validation and independent test verification show SICM6A is more accurate than the state-of-the-art methods. This, therefore, makes SICM6A provide new idea for predicting other modification sites of RNA sequences. The prediction software SICM6A is on github (https://github.com/lwzyb/SICM6A).

## INTRODUCTION

N6-methyladenosine (m^6^A) is a methylation modification of the sixth nitrogen of the post-transcriptional adenosine base^1^. As the studies have shown, m^6^A modification involves a variety of biological activities, such as mRNA splicing and stabilization, protein translation and localization, stem cell differentiation and reprogramming, biological clock, cerebellar development, and tumorigenesis^2–4^. The m^6^A site is evolutionarily conserved and has the consensus topology: DRACH motif for ***Mus musculus*** and ***Homo sapiens***^5^ (where, D = A, G or U; R = A or G; and H = A, C or U), RRACH motif^6^ for ***Arabidopsis thaliana*** and RGAC motif^7^ for ***Saccharomyces cerevisiae***. In the large range genome, methylated RNA immunoprecipitation sequencing techniques such as MeRIP-seq or m6A-seq can identify m^6^A sites in many species^8, 9^. Thus, in the whole genome analysis, high-throughput sequencing and wet experiments are the most expensive biochemical methods, hence the relatively high positive rate of false site identification^10^. Therefore, the use of computational tools to establish accurate models for predicting m^6^A sites is essential.

Current studies indicate that it is feasible to predict m^6^A sites from RNA sequences. The ***Saccharomyces cerevisiae*** methods of m^6^A site modification such as iRNA-Methyl^11^, iDNA6mA-PseKNC^12^, TargetM6A^13^, RNA-MethylPred^14^, M6Apred^15^ appeared earlier. M6APred-EL^16^ is the recent way to predict M6A sites of ***Saccharomyces cerevisiae***. RFATHM6A^17^ is a tool for predicting m^6^A sites in ***Arabidopsis thaliana***. The tools encode RNA sequences using physical and chemical properties, motifs, and space structural features. The traditional machine learning techniques such as support vector and random forest are some of the methods used for classification or prediction. The feature extraction process of these methods is very complex, and the training time is too long normally. Therefore, these methods are not suitable for large data sets, for example, ***Mammalia*** data sets of SRAMP. SRAMP^18^ also has improved traditional machine learning methods to predict m^6^A sites of ***Mammalia*** by combining the scores of several random forest classifiers. If the RNA secondary structure is analyzed before SRAMP prediction, and the calculation amount is very large, so the feature extraction time is very long. WHISTLE uses support vector machines to predict N6-methyladenosine sites in human epithelial cells, and provides query service for m^6^A sites^19^. BERMP^20^ introduced deep learning BGRU net to the prediction of m^6^A sites on SRAMP data sets for the first time, the calculation time can be reduced and the deep learning model shows a particular application prospect. Its prediction accuracy of m^6^A sites is higher in ***Arabidopsis*** and ***Mammalia*** datasets, but lower for ***Saccharomyces cerevisiae***. Therefore, only the method combined the scores between random forest and GRU can help more precisely predict the m^6^A sites of ***Saccharomyces cerevisiae***. DeepM6ASeq^21^ applies one-dimensional convolution layers to m^6^A site prediction of ***Mammalia***, the ***Mammalia*** data set for SRAMP has had much removal of redundancy. Compared with traditional machine learning, the existing deep learning classifiers are superior for species with large data sets of ***Mammalia***, but inferior to those with small data sets of ***Saccharomyces cerevisiae.*** It means that the compatibility of existing deep learning models in different species remains a problem, and the deep learning models have not been fully applied to the prediction of m^6^A sites across species. Therefore, this paper has constructed a deep learning model named SICM6A (Figure 1) for predicting m^6^A sites across species.

**Figure 1.**
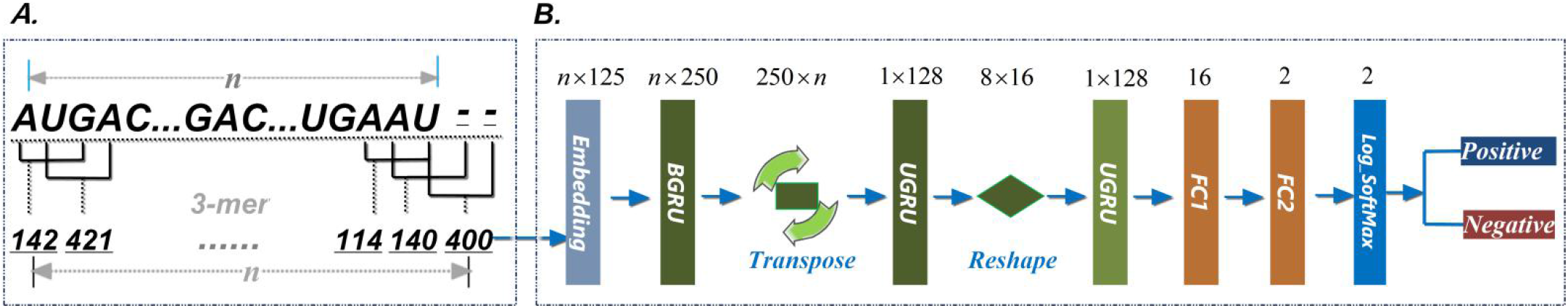
SICM6A model. ***A.*** *3-mer* encoding. The string-, A, G, C and U are represented of 5 symbols: 0, 1, 2, 3, 4 respectively, then AUG->142. Each *3-mer* nucleotides, are encoded from left to right on the sequence fragment, and the sequence fragment changes to a short vector. ***B***. SICM6A net. The word-vector embedding layer converts the short vector into a two-dimensional vector (*n×125*). The two-dimensional vector are input into a transposed and reshape GRU structure---three GRU layers in turn. The dimensional transpose operation in the middle of the front two GRU layer and the dimensional reshape operation in the middle of the last two UGRU layers. Then *1×128* dimensional matrix is output to the fully connected layer FC1 and FC2. Next, the score of the sequence fragment is computed by the log_Softmax layer. At last, a threshold of score is applied to segment, the sequence fragment is classified as positive while the score is greater than the threshold, otherwise is negative.

## METHODS

The datasets of SICM6A are from the BERMP data sets and DeepM6Aseq. The BERMP data sets are sorted from SRAMP data sets and divided into three types: ***Saccharomyces cerevisiae, Arabidopsis thaliana***, and ***Mammalia***, wherein the ***Mammalia*** data set is included the full transcription and mature mRNA. The BERMP data set for each species includes a training set and an independent test set. The DeepM6Aseq datasets include a mammalian training set that is extracted from SRAMP and an independent test set that is retrieved from the real peak data of HepG2 and human brain cell line. This paper adopts BERMP datasets to test the cross-species prediction ability of SICM6A model and DeepM6Aseq datasets to verify the identification accuracy of experimental data.

Sequence fragments are extracted from every mRNA sequence of the data sets, where DRACH, RGAC or GAC motif is in the middle. The window length *n* of a sequence fragment is 51 bits for ***Saccharomyces cerevisiae***, 101 bits for ***Arabidopsis thaliana*** and HepG2-the human brain, 251 bits for ***Mammalia*** mature RNA and 501 bits for ***Mammalia*** full transcription. Next, SICM6A uses *3-mer* encoding (Figure 1.A) to encode the sequence fragments. Each *3* consecutive nucleotides are extracted from left to right in turn, and are encoded as integer vector. The gap symbol “-” and the four types of nucleotides, A, G, C and U are commonly composed of 5 symbols: 0, 1, 2, 3, 4. *3-mer* nucleotides are converted to three numbers that combined as a integer. For example, as U->4, A->1, G->2, then *3-mer* string UAG->412. After that, *3-mer*nucleotides are encoded one by one from left to right on the sequence fragment, so each *3-mer* nucleotides are therefore encoded as an integer and the sequence fragment changes to a short vector.

The SICM6A net what it named the transposed GRU network (Figure 1.B) analyses the short integer vector. SICM6A uses the word-vector embedding layer to convert the short vector into a two-dimensional matrix (*nx125*). The features of the two-dimensional matrix are processed in turn by three GRU layers: BGRU, UGRU and UGRU. The num_embeddings and embedding_dim of the word-vector embedding layer are 1024, 125 separately. The first GRU layer’s parameters like as: the number of layers is 2, input_size is 125, hidden_size is 125, dropout is 0.5 and bidirectional is equal to “True”. The second GRU layer has 2 layers, input_size is *n*, hidden_size is 128, dropout is 0.5. The third GRU layer has 2 layers, input_size is 16, hidden_size is 128, dropout is 0.5. The dimensional transpose operation is in the middle of the front two GRU layer and the dimensional reshape operation is in the middle of the last two UGRU layers. The BGRU layer outputs *nx250* dimensional matrix, next the matrix is transposed and the UGRU generates *1×128* dimensional matrix. After the matrix is reshaped to *8×16* dimension, the third GRU layer outputs *1×128* dimensional matrix to the fully connected layer FC1 (Output=16) and FC2 (Output=2). The three layers of LeakyReLU (negative_slope=0.2), BatchNorm1d (momentum=0.5) and Dropout(p=0.25) are in the middle of FC1 and FC2. Finally, the log_Softmax layer computes the score of the sequence fragment. The sequence fragment is positive while the score is greater than a threshold, otherwise is negative. The thresholds are generally determined based on the scores of the independent test sets, corresponding to three grades of high (95%), medium (90%), low (85%) specificity (***Sp***).

The SICM6A model adopts Adam optimizer, with the learning rate of 0.0001 and the CrossEntropyLoss. The batch numbers of full transcription, mature mRNA, ***Arabidopsis thaliana***, ***Saccharomyces cerevisiae*** and HepG2_human brain cell line are 400, 1000, 200, 200, 85 respectively. The model was trained by the Pytorch framework with the mixed precision training plugin “apex” using the GPU card GTX980TI 6G. The opt_level of apex is “O2” for SICM6A. To optimize Pytorch, apex uses FP16 to speed up tensor calculation, FP32 to calculate loss value, and completes loss scaling by multiplying or dividing the scaling factor. Therefore, a small amount of GPU RAM can improve the speed and performance of the model.

## RESULTS

This paper performed five cross-validations on each ***Mammalia*** training data set and ten cross-validations on ***Saccharomyces cerevisiae, Arabidopsis*** training data sets respectively. The AUC (Area Under Curve) values of cross-validation are 0.9080, 0.8314, 0.9376, 0.8035 for full transcription, mature mRNA, ***Arabidopsis thaliana and Saccharomyces cerevisiae*** (Figure 2.a). In addition, the four models were also trained to predict the four corresponding independent test data sets. The AUC values are 0.9122, 0.8323, 0.9433, 0.8170 on the four types of independent test data sets (Figure 2.b). The results of cross-validation and independent test have similar AUC values. The AUC of SICM6A on independent test data sets is higher than state-of-the-art methods like BERMP, DeepM6A, SRAMP and M6APred-EL etc. too (Table 1). The figures indicate that the SICM6A model has better robustness and can be feasible for predicting m^6^A sites across species.

**Figure 2.**
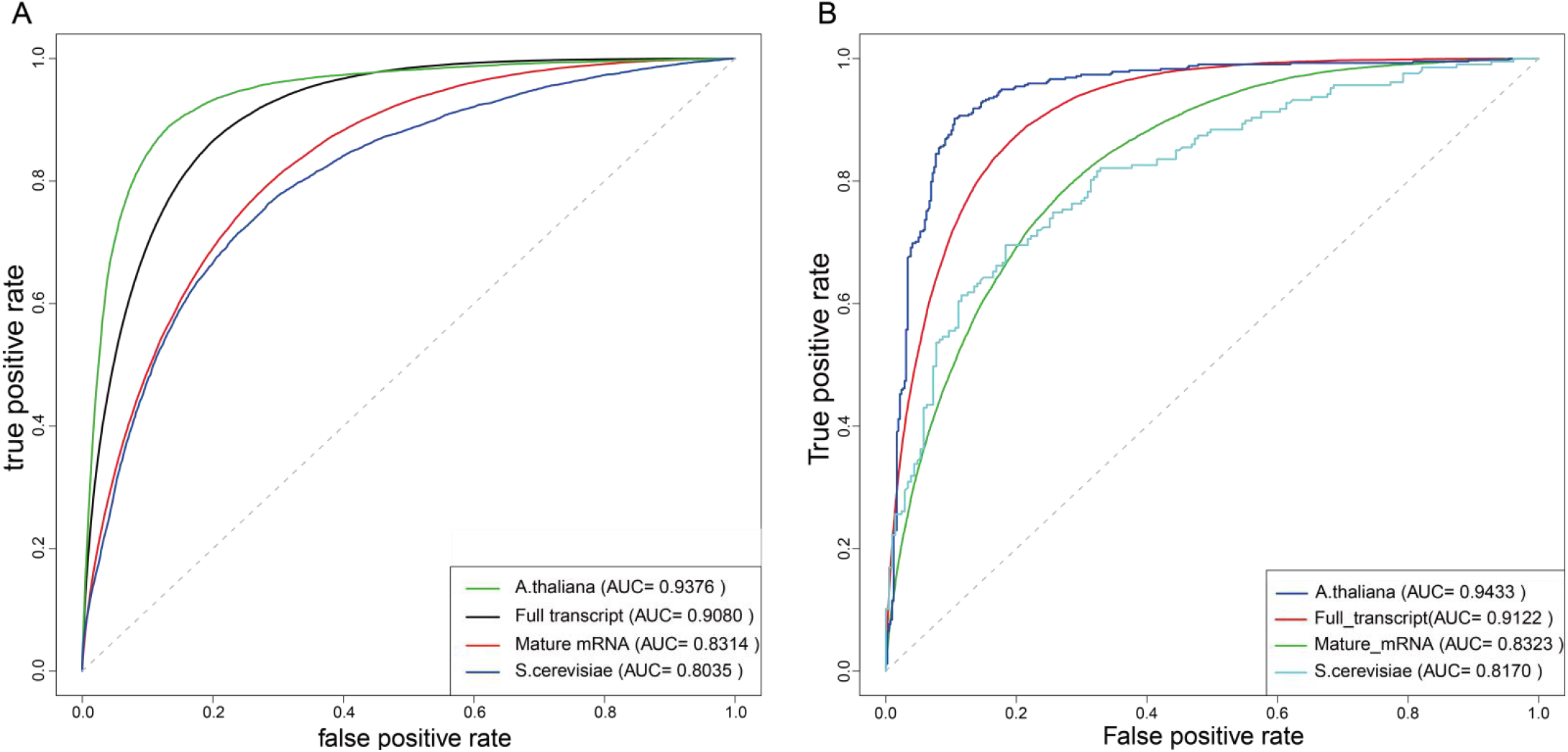
Performance of SICM6A. ***A***. Cross-validation Performance of SICM6A on the training data sets. Five cross validation is used to full transcription and mature mRNA data sets, and ten cross validation to ***Saccharomyces cerevisiae, Arabidopsis thaliana*** data sets. ***B***. Prediction performance of SICM6A on the independent testing data sets.

**Table 1.** Prediction results of different classifiers on independent datasets.

| Species | Classifiers | *Acc* | *Sn* | *Sp* | *MCC* | *AUC* |
| --- | --- | --- | --- | --- | --- | --- |
| ***Mammalia*** Full transcript | SICM6A<sup>1</sup> | 88.30 | **71.22** | 90.00 | **48.5** | **91.2** |
|  | BERMP<sup>2</sup> | 88.34 | 70.72 | 90.10 | 48.4 | 90.7 |
|  | DeepM6ASeq<sup>3</sup> | 85.81 | 44.14 | 89.98 | 29.0 | 82.9 |
|  | SRAMP<sup>4</sup> | - | 64.50 | 88.10 | 40.5 | 87.1 |
| ***Mammalia*** Mature mRNA | SICM6A<sup>1</sup> | **86.33** | 48.86 | 90.01 | 32.8 | **83.2** |
|  | BERMP<sup>2</sup> | 86.32 | 49.42 | 90.00 | 33.1 | 82.9 |
|  | DeepM6ASeq<sup>3</sup> | 84.77 | 32.39 | 89.99 | 19.8 | 73.3 |
|  | SRAMP<sup>4</sup> | - | 44.00 | 90.00 | 29.4 | 79.4 |
| ***Arabidopsis thaliana*** mRNA | SICM6A<sup>1</sup> | **89.11** | **88.28** | 89.95 | **78.2** | **94.3** |
|  | BERMP<sup>2</sup> | 87.20 | 84.21 | 90.19 | 74.5 | 93.4 |
|  | DeepM6ASeq<sup>3</sup> | 87.80 | 85.65 | 89.95 | 75.7 | 93.0 |
|  | RFAthM6A<sup>5</sup> | - | 82.20 | 90.00 | 72.5 | 93.6 |
| ***Saccharomyces cerevisiae*** mRNA | SICM6A<sup>1</sup> | **72.71** | **55.56** | 89.86 | **48.3** | **81.7** |
|  | BERMP<sup>2</sup> | 68.84 | 47.34 | 90.34 | 41.7 | 80.0 |
|  | DeepM6ASeq<sup>3</sup> | 70.29 | 50.72 | 89.85 | 44.1 | 79.7 |
|  | M6APred-EL<sup>6</sup> | 71.26 | 52.66 | 89.86 | 45.8 | 80.8 |
|  | pRNA<sub>m</sub>-PC<sup>7</sup> | 69.74 | 69.72 | 69.75 | 40.0 | 76.2 |
|  | M6A-HPCS<sup>8</sup> | 67.33 | 71.77 | 69.75 | 35.0 | 71.3 |
|  | RAM-NPPS<sup>9</sup> | 70.77 | 72.46 | 69.08 | 42.0 | 78.0 |

Note: As it is hard to get the codes of BERMP, SRAMP, RFAthM6A, M6APred-EL, pRNA<sub>m</sub>-PC, M6A-HPCS and RAM-NPPS, the comparison data is excerpted from their papers. 1. The data sets come from BERMP<sup>20</sup>. 2. The data is excerpted from BERMP<sup>20</sup>. 3. DeepM6ASeq is re-implemented using the data sets same as SICM6A. 4. The data is excerpted from SRAMP<sup>18</sup>. 5. The data is excerpted from RFAthM6A<sup>17</sup>. 6. M6APred-EL is re-implemented using the data sets same as SICM6A. 7. The data is excerpted from pRNA<sub>m</sub>-PC<sup>22</sup> and BERMP<sup>20</sup>. 8. The data is excerpted from M6A-HPCS<sup>23</sup> and BERMP<sup>20</sup>. 9. The data excerpted from BERMP<sup>20</sup>.

By comparing with advanced methods, SICM6A exhibits excellent predictive performance too. The ***Acc*** (Accuracy), ***Sn*** (Sensitivity), ***Sp*** (Specificity), ***MCC*** (Matthews Correlation Coefficient) of SICM6A are: 72.71, 55.56, 89.86, 48.3, 81.7 on ***Saccharomyces cerevisiae*** data set; 89.11, 88.28, 89.95, 78.2 on ***Arabidopsis thaliana*** data set; 86.33, 48.86, 90.01, 32.8 on mature mRNA data set; 88.30, 71.22, 90.00, 48.5 on full transcription (Table 1). The metric data ***Acc***, ***Sn***, ***Sp***, ***MCC*** are better than existing methods on ***Saccharomyces cerevisiae*** and ***Arabidopsis thaliana*** data sets. The ***Acc***, ***Sn***, ***Sp***, ***MCC*** of SICM6A are greater than SRAMP and DeepM6A on mature mRNA and full transcription data sets. The ***Sn***, ***MCC*** of SICM6A are better than BERMP on full transcription data set and ***Acc*** is greater on mature mRNA data set, other metric values are close to BERMP. The metrics show that the SICM6A model has good stability not only for a small amount of data, ***Saccharomyces cerevisiae***, and ***Arabidopsis thaliana***, but also for ***Mammalia*** with a large amount of data. Meanwhile, the ***Acc, F1-score, MCC, AUC*** of SICM6A are 78.0, 0.828, 52.6, 83.21 (Table 2) on the HepG2 and human brain cell line data sets, and greater than DeepM6ASeq and SRAMP. It represents SICM6A could identify m^6^A sites from the experimental data.

**Table 2.** Performance comparison on the HepG2 and human brain cell line data set ^1^.

| Classifiers | *Acc* | *F1-score* | *MCC* | *AUC* |
| --- | --- | --- | --- | --- |
| SICM6A | **78.0** | 0.828 | **52.6** | **83.21** |
| DeepM6ASeq<sup>2</sup> | 76.3 | 0.808 | 49.9 | 82.80 |
| SRAMP-Mature<sup>2</sup> | 73.2 | 0.787 | 42.8 | 78.50 |
| SRAMP-Full<sup>2</sup> | 76.2 | 0.824 | 48.3 | 81.80 |

Note: 1. The training and independent test data sets comes from DeepM6ASeq<sup>21</sup>. 2. The metric data is excerpted from DeepM6ASeq<sup>21</sup>.

## CONCLUSIONS

All in all, a universal classifier can help identify m^6^A sites of multiple species, so it is important to establish a predictive model with such a function. However, the current state of research is that m^6^A site prediction methods of ***Saccharomyces cerevisiae*** are often unable to accurately predict mammalian m^6^A sites, and vice versa^18^. In view of this situation, this paper made innovations in coding, model and training methods, which were different from traditional methods, and constructed the prediction method of m^6^A sites based on deep learning, SICM6A. In terms of coding, this paper adopted a very efficient *3-mer* coding. Compared with complex traditional coding, the coding method of SICM6A was not only very simple and intuitive, but also did not spend a lot of time on feature coding. On the model side, SICM6A used the word vector layer to learn the *3-mer* encoding value to find its best feature vector. In the processing of the feature vector, the SICM6A introduced the transpose operation of feature into the middle of the first two GRU layers, namely BGRU layer and UGRU layer. This novel approach is similar to extracting features by horizontal and vertical filtering, the first GRU layer could capture features from the mammalian large data set, and the second GRU layer could capture features from the ***Saccharomyces cerevisiae*** small data set. In training, in order to solve the bottleneck problem of training large data sets on medium and low level GPU graphics cards - time consuming, this paper used the mixed precision training method, which greatly accelerated the training speed and improved the prediction accuracy. Therefore, through the above improved methods, this paper subtly solved the cross-species prediction problem of deep learning, so that the SICM6A model can quickly capture the m^6^A site features from large and small data sets.

Lastly, to facilitate the identification of m^6^A sites by biologists, it developed the pytorch based software SICM6A (Figure 3), which integrates the deep learning model SICM6A proposed in this paper. The source codes are on github (https://github.com/lwzyb/SICM6A).

**Figure 3.**
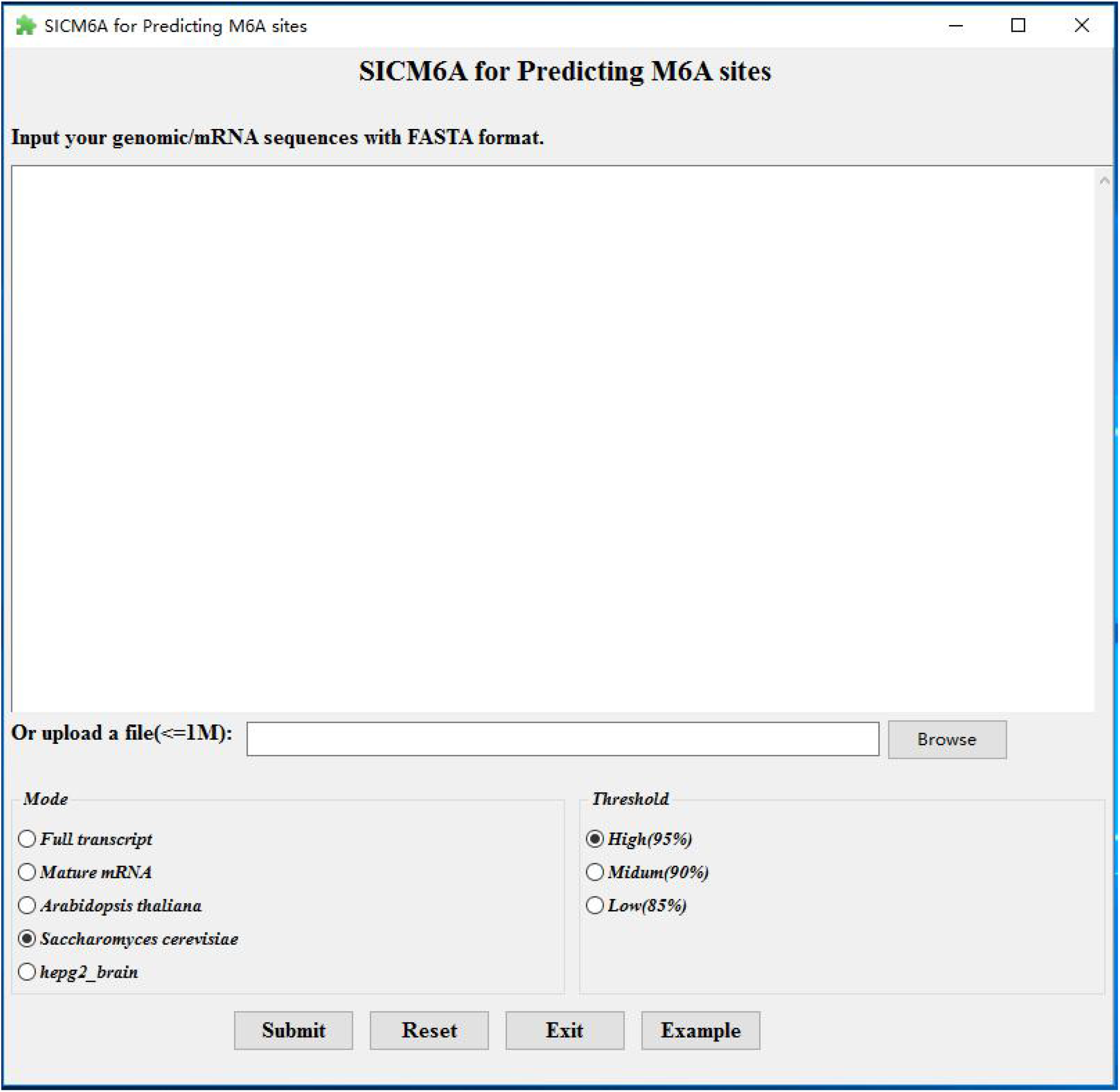
Interface of SICM6A.The user can paste the sequence into the text box and load the sequence text file in fasta format. Users can choose four species of full transcription, mature mRNA, ***Arabidopsis thaliana*** and ***Saccharomyces cerevisiae***, and set three thresholds of high, medium and low. After the user submits, the software generates the scores of short sequence fragments.

## Notes

#### Summary of Updates

Revised the introduction on the prediction tools in the introduction section

## REFERENCE

1. Liu, J. & Jia, G. Methylation Modifications in Eukaryotic Messenger RNA. Journal of Genetics & Genomics 41, 21–33 (2014).

2. Karikó, K., Buckstein, M., Ni, H. & Weissman, D. Suppression of RNA Recognition by Toll-like Receptors: The Impact of Nucleoside Modification and the Evolutionary Origin of RNA. Immunity 23, 165–175 (2005).

3. Jia, G. et al. N6-methyladenosine in nuclear RNA is a major substrate of the obesity-associated FTO. Nature Chemical Biology 7, 885–887 (2011).

4. Nilsen, T.W. Molecular biology. Internal mRNA methylation finally finds functions. Science 343, 1207–1208 (2014).

5. Dan, D. et al. Topology of the human and mouse m6A RNA methylomes revealed by m6A-seq. Nature 485, 201 (2012).

6. Levis, R. & Penman, S. 5′-Terminal structures of poly(A)^+^ cytoplasmic messenger RNA and of poly(A)^+^ and poly(A)− heterogeneous nuclear RNA of cells of the dipteran Drosophila melanogaster. Journal of Molecular Biology 120, 487–515 (1978).

7. Schraga, S. et al. High-resolution mapping reveals a conserved, widespread, dynamic mRNA methylation program in yeast meiosis. Cell 155, 1409–1421 (2013).

8. Meyer, K. et al. Comprehensive Analysis of mRNA Methylation Reveals Enrichment in 3′ UTRs and near Stop Codons. Cell 149, 1635–1646 (2012).

9. Dan, D., Sharon, M.M., Mali, S.D., Ninette, A. & Gideon, R. Transcriptome-wide mapping of N(6)-methyladenosine by m(6)A-seq based on immunocapturing and massively parallel sequencing. Nature Protocols 8, 176–189 (2013).

10. Zhang, J., Feng, P., Lin, H. & Chen, W. Identifying RNA N6-Methyladenosine Sites inEscherichia coliGenome. Frontiers in Microbiology 9, 955- (2018).

11. Chen, W., Feng, P., Ding, H., Lin, H. & Chou, K.C. iRNA-Methyl: Identifying N 6-methyladenosine sites using pseudo nucleotide composition. Analytical Biochemistry 490, 26–33 (2015).

12. Feng, P. et al. iDNA6mA-PseKNC: Identifying DNA N 6-methyladenosine sites by incorporating nucleotide physicochemical properties into PseKNC. Genomics 111, S0888754318300090 (2018).

13. Li, G.Q., Liu, Z., Shen, H.B. & Yu, D.J. TargetM6A: Identifying N6-methyladenosine Sites from RNA Sequences via Position-Specific Nucleotide Propensities and a Support Vector Machine. IEEE Trans Nanobioscience PP, 1–1 (2016).

14. Jia, C.Z., Zhang, J.J. & Gu, W.Z. RNA-MethylPred: A High Accuracy Predictor to Identify N6-methyladenosine in RNA. Analytical Biochemistry 510, 72–75 (2016).

15. Chen, W., Hong, T., Liang, Z., Lin, H. & Zhang, L. Identification and analysis of the N6-methyladenosine in the Saccharomyces cerevisiae transcriptome. Scientific Reports 5, 13859 (2015).

16. Wei, L., Chen, H. & Su, R. M6APred-EL: A Sequence-Based Predictor for Identifying N6-methyladenosine Sites Using Ensemble Learning. Mol Ther Nucleic Acids 12, 635–644 (2018).

17. Wang, X. & Yan, R. RFAthM6A: a new tool for predicting m 6 A sites in Arabidopsis thaliana. Plant Molecular Biology 96, 327–337 (2018).

18. Zhou, Y., Zeng, P., Li, Y.-H., Zhang, Z. & Cui, Q. SRAMP: prediction of mammalian N6-methyladenosine (m6A) sites based on sequence-derived features. Nucleic Acids Res 44, e91–e91 (2016).

19. Chen, K. et al. WHISTLE: a high-accuracy map of the human N6-methyladenosine (m6A) epitranscriptome predicted using a machine learning approach. Nucleic Acids Res 47, e41–e41 (2019).

20. Huang, Y., He, N., Chen, Y., Chen, Z. & Li, L. BERMP: a cross-species classifier for predicting m(6)A sites by integrating a deep learning algorithm and a random forest approach. Int J Biol Sci 14, 1669–1677 (2018).

21. Zhang, Y. & Hamada, M. DeepM6ASeq: prediction and characterization of m6A-containing sequences using deep learning. BMC Bioinformatics 19, 524–524 (2018).

22. Liu, Z. et al. pRNAm-PC: Predicting N(6)-methyladenosine sites in RNA sequences via physical-chemical properties. Analytical Biochemistry 497, 60–67 (2016).

23. Zhang, M. et al. Improving m(6)A sites prediction with heuristic selection of nucleotide physical-chemical properties. Analytical Biochemistry 508, S0003269716301154 (2016).

